# Speciated evolution of oscillatory mass-action chemical reaction networks

**DOI:** 10.1101/2025.04.10.648132

**Authors:** Lillian T. Tatka, Lucian P. Smith, Herbert M. Sauro

**Affiliations:** Department of Bioengineering, University of Washington, Seattle, WA, USA

## Abstract

Evolutionary algorithms, a class of optimization techniques inspired by biological evolution, have emerged as powerful tools for the optimization of complex systems, including the evolution of mass-action chemical reaction networks. This work explores the application of evolutionary algorithms in this domain, presenting a novel approach inspired by neural network evolution methodologies. A key feature of the algorithm is speciation, which separates candidate reaction networks into groups based on their similarity, which maintains diversity and protects innovations. Crossover has also been shown to be an effective means of improving evolutionary success in other domains. However, crossover of mass-action networks is tested and found to be detrimental to the evolutionary process. This work goes beyond theoretical exploration by offering a practical contribution in the form of a user-friendly software module. This module encapsulates the newly devised algorithm, enabling researchers and practitioners to readily apply the speciation-based approach in their own investigations of mass-action chemical reaction networks.

**Author summary:** Evolutionary algorithms are an optimization technique inspired by biological evolution. They can be used to solve complex multi-dimensional problems for which analytic solutions are infeasible. We developed a novel evolutionary algorithm for use with mass-action chemical reaction networks. This algorithm implements two features of biological evolution, speciation and crossover, in an effort to generate chemical reaction networks with specific behaviors. Here, this novel algorithm is demonstrated by generating chemical reaction networks whose chemical species oscillate in time. This algorithm is encapsulated in a julia package and is publicly available as ReactionNetworkEvolution.jl.

## 1 Introduction

Genetic algorithms (GA), inspired by the principles of natural selection and genetics, have emerged as a powerful computational technique to solve optimization and search problems. Developed by John Holland in the 1970s, GAs mimic the process of biological evolution to solve optimization and search problems [1]. At their core, genetic algorithms operate on a population of potential solutions, represented as individuals or “chromosomes,” each encoding a candidate solution to the problem at hand. Through iterative generations, genetic algorithms apply mechanisms of selection, crossover, and mutation to evolve and refine the population, gradually converging towards optimal or near-optimal solutions. Selection favors individuals with higher fitness, mirroring the process of natural selection, while crossover and mutation introduce variation and diversity into the population, allowing exploration of the solution space. Using the principles of evolution, genetic algorithms offer a versatile and robust approach to solving complex problems in various domains, from engineering and optimization to biology and beyond.

Evolutionary algorithms offer a computational framework for addressing the inherent complexity and nonlinearity of biological systems, providing a means to explore large solution spaces and optimize model parameters and structures. The use of evolutionary algorithms in systems biology became popular in the late 1990s, often to better understand metabolic pathways. This approach has been used to demonstrate the optimality of the glycolysis [2, 3] and the Krebs citric acid cycle [4]. In these studies, genetic algorithms were used to optimize reaction rates but not the structure of the networks. Francois and Hakim evolved both the structure and reaction rates of genetic regulatory networks to generate small genetic networks that functioned as bistable switches and oscillators [5]. Likewise, Deckard and Sauro [6] used the same approach to evolve biochemical networks using a similar algorithm. In both cases the algorithm used only mutation and selection, avoiding crossover, to minimize the difference between the candidate networks’ output and idealized time series data displaying the desired behavior. Francois and Hakim argued that one of their evolved oscillators was structurally similar to the networks responsible for the circadian rhythm. However, Chu extended this work and found no consistent patterns in the evolved oscillators [7]. She noted that it is computationally unfeasible to thoroughly examine all resulting networks, but argued that the similarity to the circadian rhythm was likely coincidental. Despite Chu’s study occurring almost 20 years ago, the problem of thoroughly pattern mining evolved networks remains an issue. The results of these efforts encouraged the exploration of evolutionary techniques on similar problems.

Fujimoto et al. employed a similar strategy to study genetic networks leading to striped patterning and ultimately body segmentation in arthropods [8]. When analyzing the resulting networks, they found three distinct classes, feed-forward loops, feed-back loops, and interconnections between the two. These three classes reproduced various segmentation strategies observed in arthropods.

Kobayashi et al. also explored the evolution of oscillatory genetic networks, however their algorithm fixed parameter values and only allowed connection rewiring to achieve oscillations with prescribed time periods [9]. Deckard and Sauro [6] evolved small networks with specific computational capabilities such as square root and cube root calculators. As with the previously mentioned algorithms, their algorithm avoided crossover, relying solely on mutation to develop candidate solutions. Attempts to use crossover did not result in fitter offspring, possibly due to combining heterologous networks and thus disrupting developing solutions. This work was continued by Paladugu et al., who sought to generate mass-action network functional modules with specific behaviors, including oscillators [10], switches, and various frequency filters. This algorithm avoided crossover as the authors argued that it would be disruptive.

Marchisio and Stelling argued that brute force optimization algorithms lack efficiency and that implementing some element of rational design could speed up automatic design of genetic networks implementing boolean logic gates [11]. They separate structural and parameter optimization by first generating several possible circuits and then ranks feasibility based on the structure. Then the best solutions undergo parameter optimization. Separating structural design from parameter optimization certainly saves computation time, but it relies on the existence of a rational solution, and is better suited for genetic networks, where these attributes are more easily separated. For more complex tasks, a solution might depend on a precise combination of both structure and parameter.

Other approaches fix structure and make use of evolutionary algorithms to explore the parameter space while fixing network topology. Jin and Sendhoff used an evolution strategy to explore parameters of regulatory motifs with fixed structures [12]. Porubsky and Sauro use an evolutionary algorithm to search for parameters that yield oscillations in mass-action networks of fixed structure [13].

Most applications of evolutionary algorithms in systems biology avoid the use of crossover as it tends to disrupt candidate solutions. For example, Drennan and Beer successfully evolved a repressilator using a genetic algorithm that included crossover [14]. Instead of evolving sets of differential equations, Drennan and Beer encoded their genetic regulatory networks using strings of 4 different letters, analogous to DNA encoding. Knabe et al. used a similar approach to evolve oscillators that interact with the environment [15]. This approach allows for the straightforward implementation of crossover, but limits the space of possible models. Due to this encoding, there was a high probability of breaking connections making it difficult to pass on structural innovations.

Stanley and Miikkulainen addressed this problem in 2002 with NEAT (NeuroEvolution of Augmenting Topologies), a powerful algorithm used for evolving artificial neural networks (ANNs) [16]. Unlike traditional methods of neural network training, which typically involve adjusting the weights and biases of fixed network architectures, NEAT evolves both the structure and weights of neural networks simultaneously. Two key features of NEAT are the use of a more meaningful structural crossover technique, and speciation, which protects innovations and allows them time to develop by having candidate solutions compete only against similar individuals instead of the entire population. Dinh et al. adapted this technique to genetic regulatory networks and found improved performance in evolving oscillators with the use of crossover [17].

A key feature of genetic algorithms is crossover, the exchange of multiple parameters between two candidate solutions to create a new offspring solution. Crossover proves challenging when solutions are not vectors of matrices of numbers, but also include graphs. Stanley and Miikkulainen devised a method, NEAT, to crossover topologies when evolving artificial neural networks [16]. Dinh et al. adapted NEAT to enable crossover when evolving gene regulatory networks [17]. Here, the NEAT algorithm is adapted and explored with mass-action chemical reaction networks. In contrast to previous studies in different domains, the crossover method presented here does slightly reduce the success rate of evolution.

Speciation, though not essential to evolutionary algorithms, can improve their success by protecting innovations and allowing them to develop into better solutions. This work also introduces a method for separating chemical reaction networks into species and explores the effect of speciation on evolutionary success.

The purpose of this work is twofold: (1) to create a general purpose, easily customizable, module for evolving mass-action chemical reaction networks and (2) to explore the effects of crossover, speciation, and other hyperparameters on the success rate of the algorithm. The successful implementation of this algorithm could aid systems biologists in the construction of computational models or generate ensemble models to predict behavior. The algorithm could offer numerous possible explanations for a given phenomenon or set of time series data. In cases where multiple models explain the same phenomenon, future iterations of this software may be useful in determining what data is needed to clarify the model.

Unlike previous studies, this work is specific to mass-action networks and modularizes the evolutionary process. Users can precisely tune the evolutionary algorithm to allow or disallow crossover, specify the extent of speciation, and adjust various mutation probabilities and techniques. The evolutionary algorithm is packaged in a library written in the julia programming language [18]. It is highly configurable and allows users to easily specify settings via the command line for a simple JSON file. It can generate reaction networks to match a variety of time series data beyond oscillators.

## 2 Methods

The evolutionary algorithm begins by generating a population of random networks with a set number of chemical species and reactions (Fig 1). Each network in the population is then sorted into species groups based on its topological similarity to other networks. The fitness of each individual network is evaluated and each species group is assigned the fitness of its top performing network which determines the number of offspring each species gets to contribute to the subsequent generation. Species with higher finesses are allowed to produce more offspring. The next generation is constructed by first selecting the top performing individuals from each species without modification. The rest of the subsequent generation is populated by networks that are modified by either crossover, mutation, or both. This new generation is then grouped into species based on structural similarity, and the process repeats. Each aspect of this algorithm is described in further detail below.

**Fig 1.**
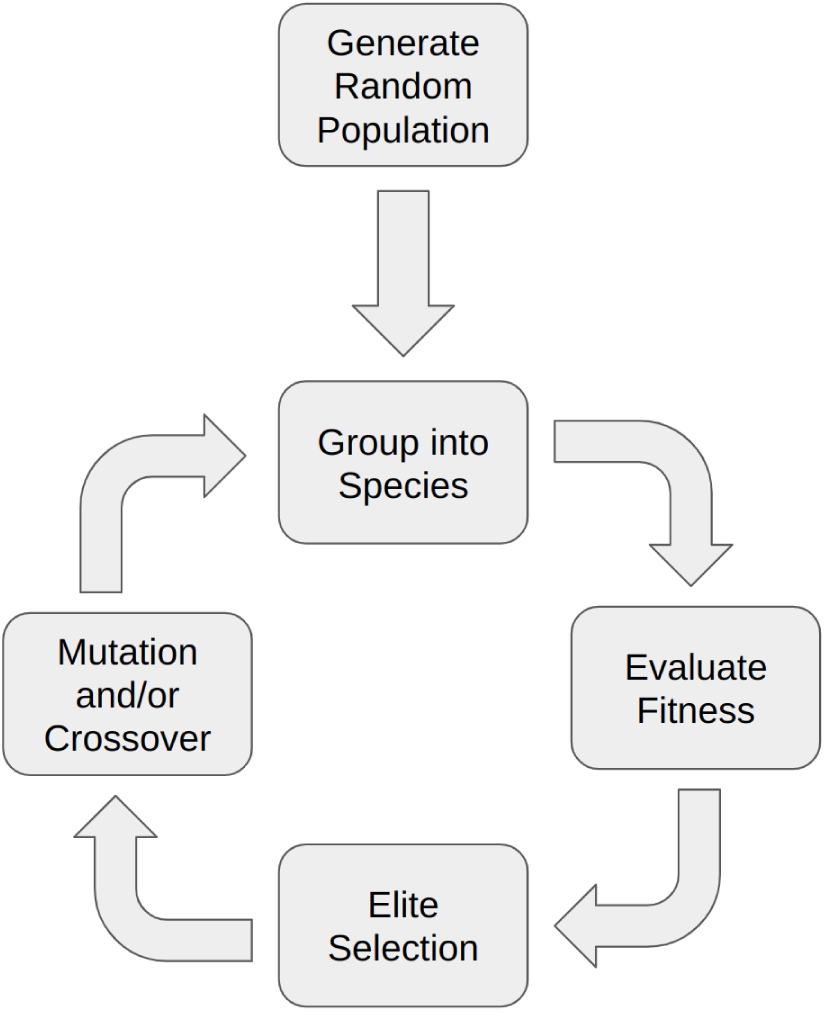
Overview of the evolutionary algorithm. The evolutionary algorithm begins with a population of randomly generated reaction networks which are then grouped into species and their fitness evaluated. Subsequent generations are populated by elite selection followed by crossover and/or mutation.

### 2.1 Encoding

Chemical reactions are represented as custom data structures that contain one or two reactants, one or two products, a rate constant, and a boolean value indicating if the reaction is active or inactive. Four types of mass-action reaction are possible: uni-uni (a single reactant to a single product), bi-uni (two reactants form a single product), uni-bi, and bi-bi. A reaction that is active participates in the network that it is part of. An inactive reaction is “turned off” but remains part of the network as a historical record. It can be activated again or crossed over under certain circumstances.

A reaction network is a custom data structure consisting of several reactions, initial concentrations for the floating species, and other information to track the individual network. The initial population of randomly generated networks had a set number of chemical species (3) and random reactions (5). The number of chemical species remained constant throughout the evolution process, but the number of reactions could vary.

### 2.2 Mutation

There are two types of mutation: rate constant mutation and reaction addition or subtraction. The probability of each of these mutations occurring and the range in which rate constants are mutated can be set by the user. A rate constant mutation can occur in two possible forms. The rate can be increased or reduce by a random percentage, uniformly distributed in a custom range (by default ±20%), or more rarely a completely new rate constant can be selected, uniformly distributed within a custom range (by default 0.1 to 50). Rate constants were not allowed to become negative, but they did not have an upper bound as previous studies observed that many synthetic oscillating systems rely on a single reaction with an abnormally large rate constant [19].

A reaction mutation is either the addition of a new random reaction or the deletion of an existing reaction, with 50% chance of each. When a new reaction is added, it is generated randomly with a probability of 0.1, 0.4, 0.4, 0.1 for uni-uni, uni-bi, bi-uni, and bi-bi reactions respectively. These probabilities are also configurable. If the randomly generated reaction is already present in the reaction network, the new reaction’s rate constant is added to the existing reaction’s rate constant. This feature can allow some rate constants to grow to several multiples of the next largest rate constant in the network. When a reaction is deleted, it is “switched off” and becomes inactive in the network, but is preserved as it can become reactivated under certain circumstances.

### 2.3 Crossover

During crossover, two parent networks from the same species group are chosen to mate. Analogous to natural evolution, crossover is essential a means of changing more parameters simultaneously and to a greater degree. It injects novelty into the offspring network compared to single mutations. This could either result in faster progress towards a good solution, or could interrupt progress by destroying promising solutions.

Crossover occurs by examining each reaction in the parent networks. If a reaction is present in both parents, the reaction will be passed down to the offspring network with the rate constant randomly chosen from one parent. If a reaction is present in the more fit parent, but not the less fit parent, the offspring network will inherit the reaction (Fig3). If an inherited reaction is currently inactive meaning it was previously deleted during a mutation step), there is a 0.25 probability of the reaction becoming active again. If both parents are equally fit, one is randomly chosen to be the more fit parent (Fig 2).

**Fig 2.**
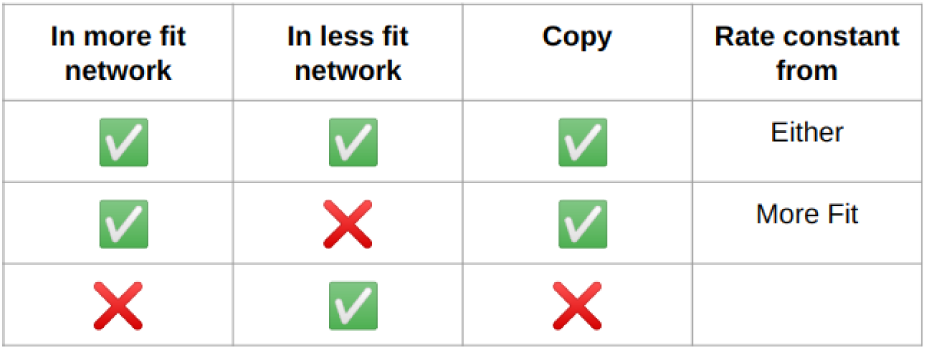
Summary of crossover procedure. Every reaction in the more fit network is passed down to the offspring network. If it occurs in both parent networks, the rate constant is randomly inherited from one parent or the other.

Figure 3 depicts the crossover of two similar parent networks and the resulting offspring. Only networks that are part of the same species group (and are thus topologically similar) can be crossed over. The structure of the more fit parent is preserved but rate constants from the less fit parent can be passed on to the offspring network.

**Fig 3.**
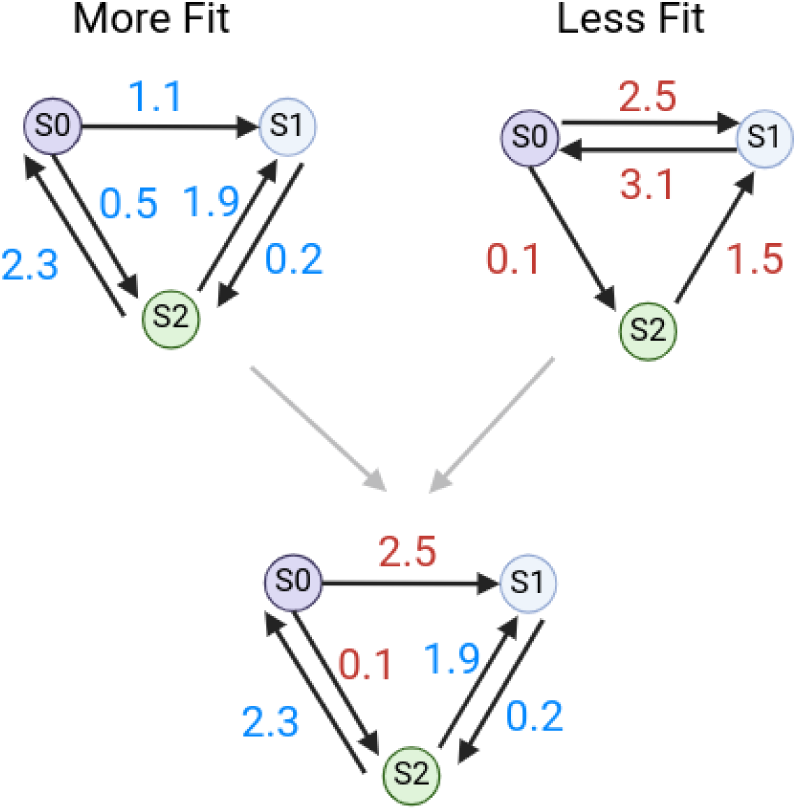
An example of network crossover. The topology of the more fit parent is passed to the offspring. Rate constants for reactions that are shared are passed down randomly.

A second crossover strategy for similar networks was tested in later trials. Networks that were members of the same species and had similar fitness scores (within 5% of each other) underwent a slightly different crossover process. All reactions that the two networks had in common were passed down to the offspring network with rate constants chosen randomly from one parent or the other. Reactions that occurred in one parent network but not the other had a 50% chance of being passed down to the offspring network.

### 2.4 Speciation

In many cases, modifying a network, especially through the addition or deletion of reactions, initially makes the system less fit. Without speciation, these innovations seldom last for more than a generation. However, these topological innovations often prove to be an essential step in evolution [16]. For this reason, a speciation strategy is employed to protect innovations and allow them time to optimize. Individuals with similar topologies are grouped into species. Members of a species compete only with each other and not the population at large. A compatibility distance metric, *δ*, is used to assign reaction networks to a species, defined as

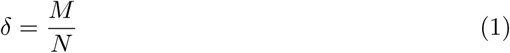

where M is the number of reactions in the larger network that are not present in the smaller network and N is the total number of reactions in the larger network. Two reactions are considered identical if they have the same reactants and products. Rate constants are not considered. If *δ* is below the speciation threshold, *δ_t_*, the two networks are members of the same species.

For each new generation, networks are assigned to species based on the previous generation’s species groups. Each new network is compared to the best network from a species in the previous generation. If *δ* is less than the speciation threshold *δ_t_*, then the network is assigned to that species. If *δ* is greater than *δ_t_*, then the network is compared to a randomly selected individual from the next species, etc., until a match is found. If no species is assigned after comparing the network to all species in the previous generation, then a new species is created.

The speciation threshold *δ_t_* is adjusted with each generation to steer the total number of species towards the target number of species. The target number of species is 10 by default, but can be configured by the user. After each generation the total number of species is counted. If there are more species than the target value, *δ_t_* is increased allowing networks that are more dissimilar to be grouped together. If there are fewer species than the target value, *δ_t_* is decreased. Adapting *δ_t_* as evolution proceeds maintains an optimal number of species. If there are too many species each with a small number of members, there are not enough opportunities to optimize solutions in a small parameter space. Similarly, if there are too few species with several members, the entire parameter space is not adequately explored.

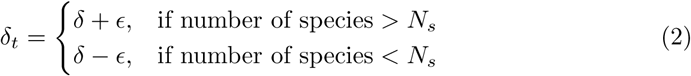

### 2.5 Objective Function

The objective function evaluates how well a candidate model oscillates at the imposed time period, *T* . Two arbitrary concentrations, [*C*]_1_ and [*C*]_2_, are chosen *apriori*. The fitness score of a network describes how well any of its chemical species approached [*C*]_1_ at half periods (*T/*2, 3*T/*2, 5*T/*2,…) and [*C*]_2_ at each period (*T*, 2*T*, 3*T*,…) over the course of 5.5 periods. The candidate model’s single chemical species that best approached these points was selected and the fitness was evaluated as the maximum of the reciprocal of the absolute difference between the time series data and [*C*]_1_ or [*C*]_2_, the “ideal oscillator” time points, (equation 3). The reciprocal was taken so that higher values represented better fitness, which is necessary for determining the number of offspring for each species in subsequent steps. This objective function has been shown to be an effective means of evolving oscillators in similar and previous work [5, 10, 19]. In cases where candidate models could not be simulated, a fitness of 0.0 was assigned, resulting in the candidate model’s subsequent removal from the population.

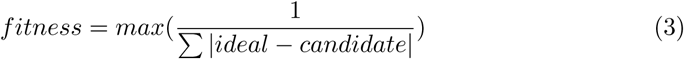

### 2.6 ODE Solver

During the fitness evaluation phase, reaction networks were converted to systems of ordinary differential equations. These equations were then numerically solved using the

DifferentialEquations.jl julia package [20] and the CVODE solver from the Sundials suite for solving initial value problems [21]. The CVODE solver was shown to be effective for this application in previous work [19].

### 2.7 Reproduction

Each model species was allocated offspring in proportion with its fitness, with more fit species producing more offspring. The fitness of species was defined as the fitness of its most fit individual. To calculate the number of offspring allocated to each species group, the total fitness of the population was calculated by summing the fitness of each species. Then each species, *s*, was assigned an offspring number proportional to its contribution to the fitness of the entire population (equation 4). By default, a single species was only allowed to produce at most 10% of the subsequent generation in order to prevent a single species from taking over the entire population (but this value could be adjusted by the user). For these reasons, the total number of individuals fluctuated slightly over time.

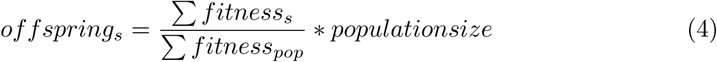

The reproduction phase began with an elitism selection strategy. The top 10% of networks in each species were copied to the next generation directly. If a species had 10 or fewer individuals, the single best network was copied without modification to the next generation. Then, the bottom 10% of networks in each species are deleted. For the remainder of the allocated offspring for each species, a random number, *p*, was generated. If the *p* was less than the probability of crossover, then two networks from the same species were selected and crossed over to produce a single offspring network. Otherwise a single network was chosen. If *p* was greater than or equal to 1 - the probability of mutation, the network was mutated. Outside of the elite networks, all networks were either crossed over or mutated. Some were both crossed over and mutated. The probabilities of crossover and mutation could be configured by the user. Probabilities for specific types of mutations are described in section 2.2.

Tournament selection was also explored and can be configured in settings. In this case, two networks from the same species were randomly chosen and the more fit network was then crossed over, mutated, or both. If crossed over, a second network was also chosen by tournament selection (figure 4).

**Fig 4.**
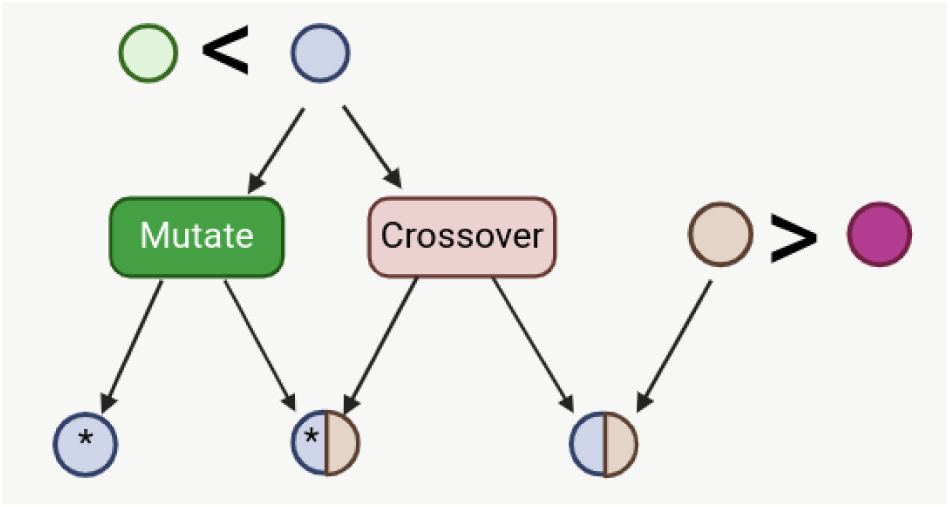
Tournament selection with mutation and/or crossover. Two networks are selected at random and the more fit network of the two undergoes either crossover, mutation, or both. If the network undergoes crossover, a mate network is also chosen by tournament selection.

### 2.8 Output

The evolutionary algorithm was repeated for a preset number of generations (800 by default) or until the best model reached a predefined fitness level. At this point, the best model from each species was converted to Antimony, a human readable model definition language [22] and written to a text file along with the model’s fitness. All settings were also written out to a JSON file, which could be used without modification to repeat the evolution trial exactly.

The best models from each evolution trial were automatically assessed with a Python script using the Tellurium modeling environment [23] and RoadRunner simulation software [24]. A model was considered an oscillator if its eigenvalues at steady state contained a positive real number with a nonzero imaginary number and if all concentrations were positive at steady state. Models that met the eigenvalue criteria but had negative steady state concentrations were flagged. Flagged models then underwent a simple automated repair process wherein each reaction was deleted one at a time in an effort to meet the oscillation criteria. Models that met criteria after the simple repair process were counted as oscillators.

Each evolution trial was run on a single CPU in a linux environment. Often several trials were run in parallel across several CPUs using the cluster computer at the University of Washington.

### 2.9 Success Metrics

A single evolution trial was considered successful if it produced a network that oscillated. For each experiment, 700 trials of evolution were performed and the total number of resulting oscillators evaluated. The success rate is the portion of the 700 models that resulted in an oscillator. Experiments sought to maximize this number. Computation time was considered qualitatively in the discussion of results. Often experiments with higher success rates came at the expense of computation time. This metric also did not consider the diversity of the networks produced. For example, a batch of several evolution trials could produce duplicate oscillating networks without penalty.

## 3 Results

The software was evaluated with multiple hyperparameter settings to assess its ability to generate oscillatory mass-action chemical reaction networks. Each set of hyperparameters were evaluated over 700 evolutionary trials and the success rate recorded. A trial was deemed successful if it generated a network with sustained oscillations, as evaluated by the criteria described previously.

### 3.1 Verifying Speciation

Before evaluating the effects of speciation on evolution, tests were run to verify that speciation was occurring and that the adaptive *δ* threshold was maintaining the target number of species. Batches of 700 trials were run for a target number of 1 (no speciation), 5, 10, 30, and 50 species. Figure 5 shows that on average, these target numbers were achieved. During the initial generations, the number of species fluctuates more dramatically. This is like due to the fact that while *δ* is adaptive, *ɛ*, the amount by which *δ* is changed, is not. During the first several generations, adjustments to *δ* tend to overshoot or undershoot the target number of species, likely due to there being large differences between individual networks at first. This over and undershooting causes the number of species to fluctuate before leveling out to the target level after approximately 15 generations.

**Fig 5.**
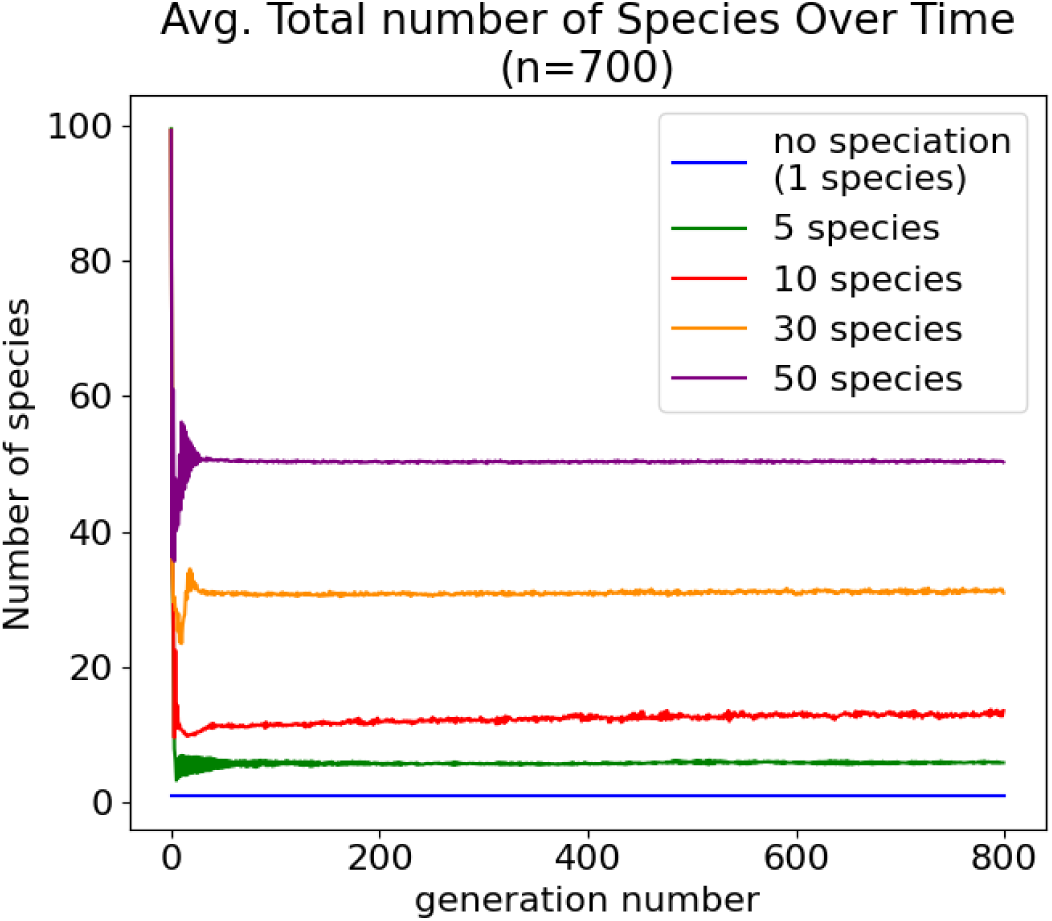
Adapting *δ* maintains the target number of species without the need to select a value that results in the target number of species through trial and error.

The benefit of an adaptive *δ* is that the user need only specify the target number of species and *δ* will change to accommodate that specification. Without adaptation, *δ* must be chosen through trial and error so that it results in the target number of species. For example, in the case of a target species of 10, with an adaptive *δ*, the total number of species converges to 10 after a few generations and remains there until the end of the trial (Figure 6). With a constant *δ*, the total number of species is gradually reduced over the course of the trial. In this example, in both cases the success rate was about the same, 22% vs 22.5% for adaptive and constant *δ*, respectively.

**Fig 6.**
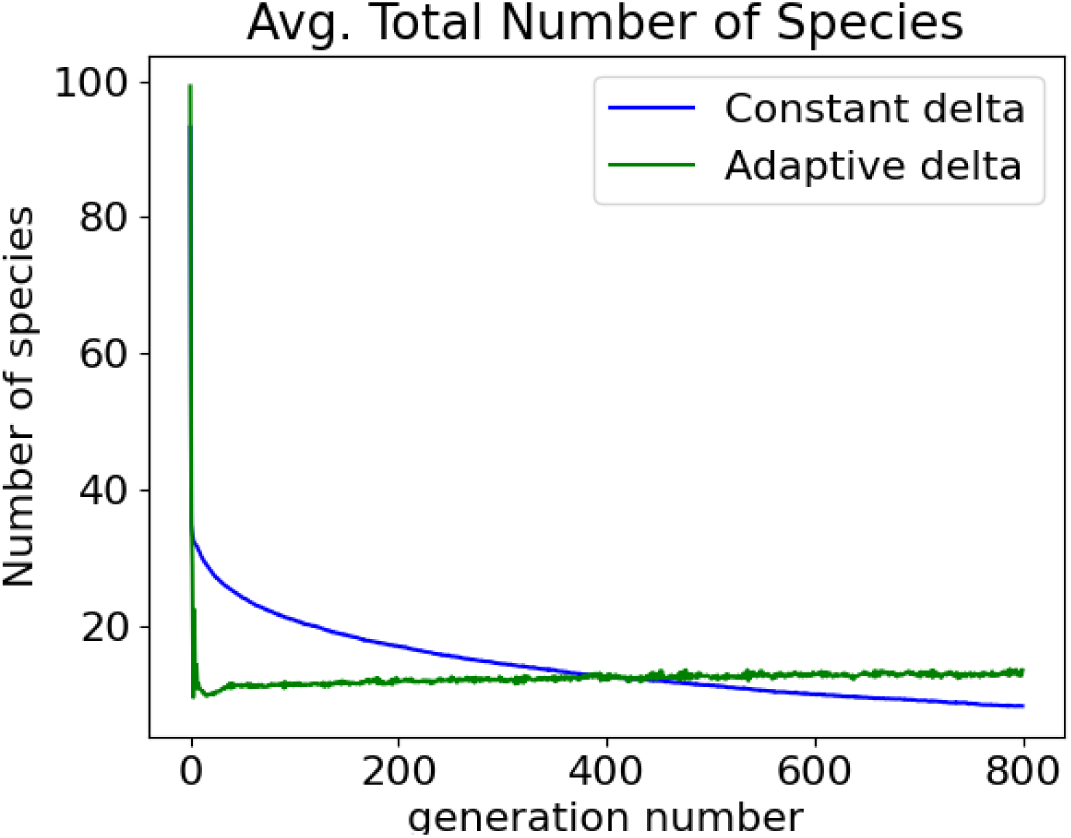
Adaptive *δ* maintains the target number of species compared to a constant delta.

### 3.2 Speciation Effects

In order to test the effects of speciation on population diversity and oscillator success rate, evolution was run without speciation, and with 10 and 30 as the target number of species. When speciation is disabled, the top 10% of the entire population is passed on to the next generation unmodified and the bottom 10% of the entire population is deleted. Then mutation is performed on the entire population to populate the remainder of the subsequent generation (crossover was not performed for these experiments). Each network must compete against every other network in the population. To account for fluctuations in population size when speciation is enabled, diversity was measured as the number of unique networks over the total number of networks in the population. A network was considered unique if differed by one or more reactions from all other networks. Reactions are considered different if they differ in product or reactant. Rate constants are not considered in these comparisons. Significance levels were calculated using two-tailed t-tests.

In trials where no speciation was allowed, the average portion of unique networks was 0.294 ± 0.035 across 800 generations. This is significantly (p *<*0.0001) less than the portion of unique networks in evolution trials with a target number of species of 10, 0.344 ± 0.037 (figure 7).

**Fig 7.**
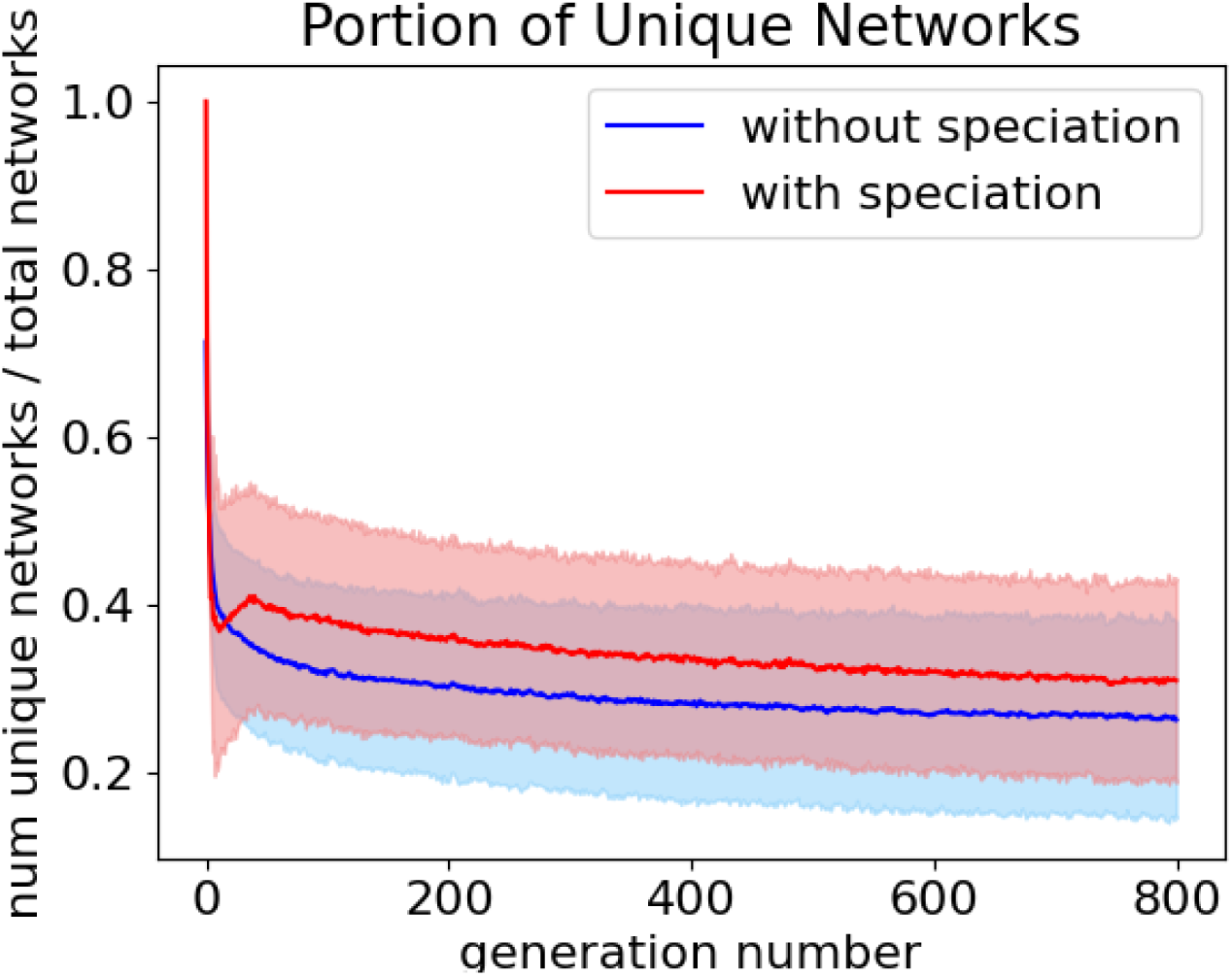
The average portion of unique networks over 800 generations with speciation (red), 0.344 ± 0.037, and without (blue), 0.296 ± 0.034. Shading is the 95% confidence interval.

One purpose of speciation was to prevent the most fit network from overtaking the population over time. When a single network dominates the population, there are not enough unique networks to adequately explore the solution space and the evolution algorithm converges on a solution prematurely. To account for fluctuating population sizes when speciation is allowed, the number of copies of the best network divided by the total number of networks was measured at each generation with speciation (with a target species number of 10) and without speciation. When speciated was enabled, the portion of the population occupied by copies of the most fit network remained constant with an average portion of best networks of 0.071 ± 0.008. Without speciation, the portion of best networks rose gradually over time with an average portion of 0.402 ± 0.079 (Figure 8). This shows that speciation effectively prevents a single network from dominating the population (p *<*0.0001).

**Fig 8.**
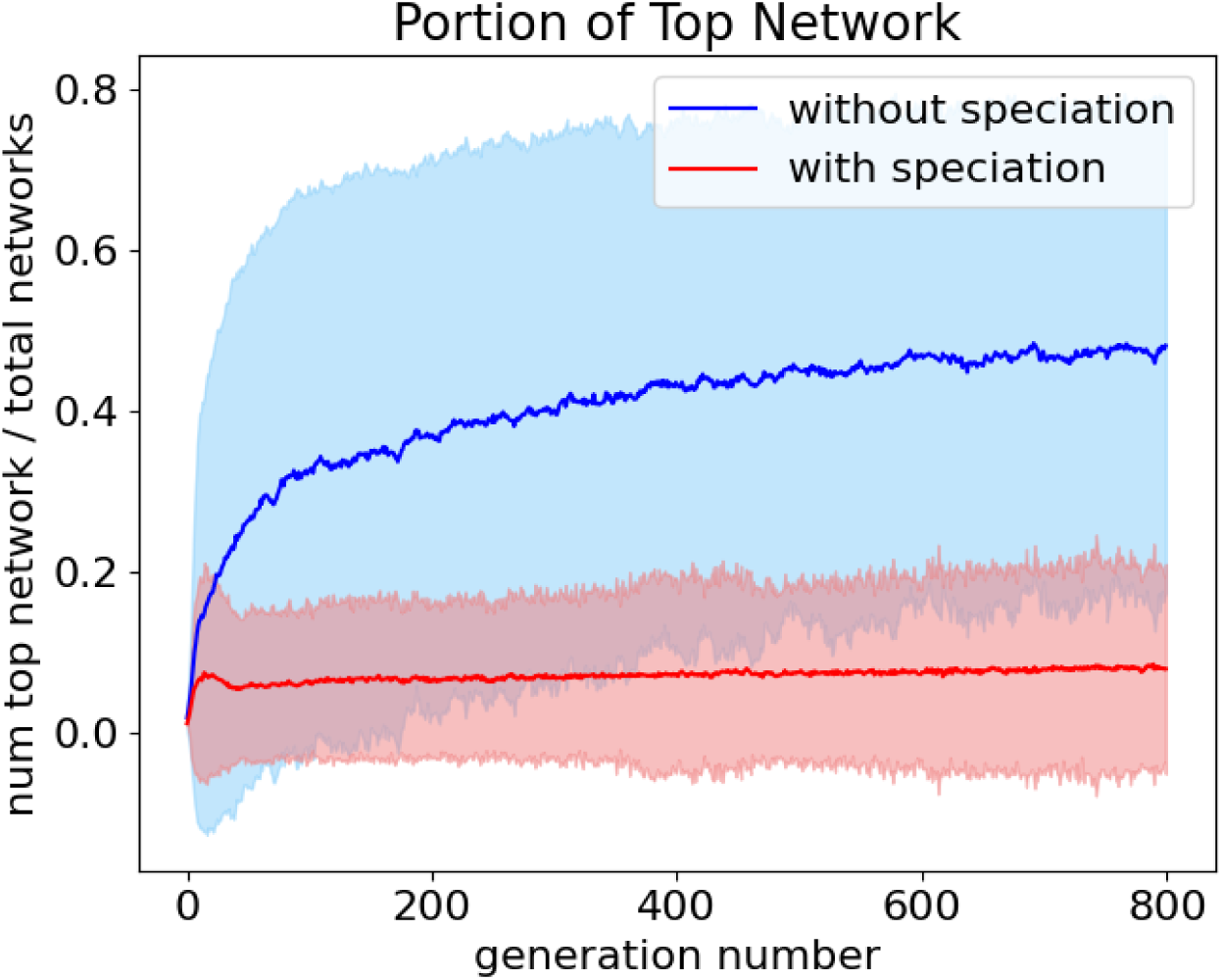
The average portion of best networks over 800 generations with speciation (red), 0.071 ± 0.008, and without (blue), 0.402 ± 0.079. Shading is the 95% confidence interval.

Next, the effects of speciation on evolution success rate was evaluated comparing no speciation to speciation with a target species number of 5, 10, 30, and 50 (Figure 9. Again, crossover was omitted from these studies. For each speciation level, a batch of 700 trials of evolution were run and the number of oscillators resulting from each batch. Significance levels were calculated using a binomial test.

**Fig 9.**
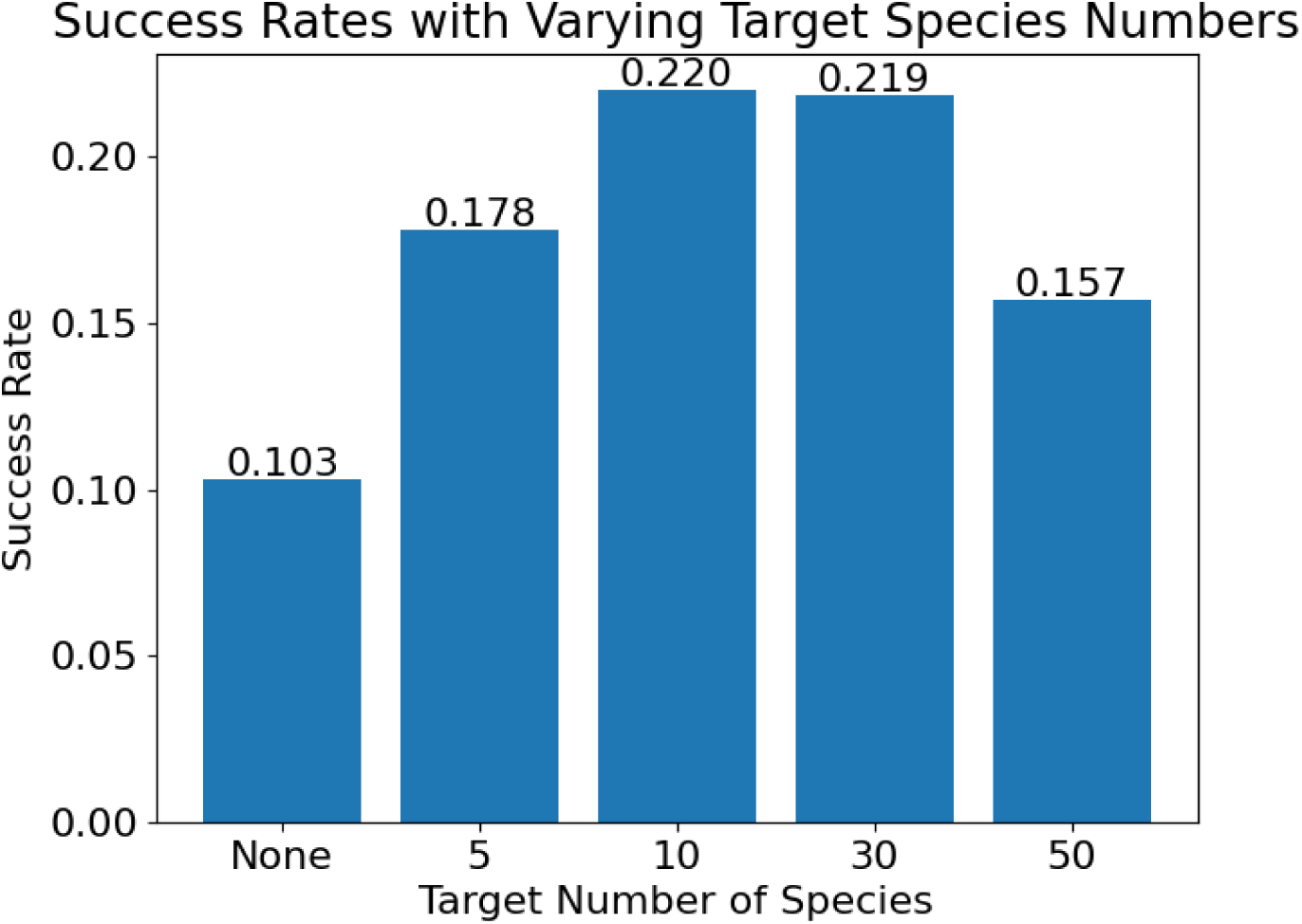
Evolution success rate of 700 trials with no speciation and with a target species of 5, 10, 30, and 50.

**Fig 10.**
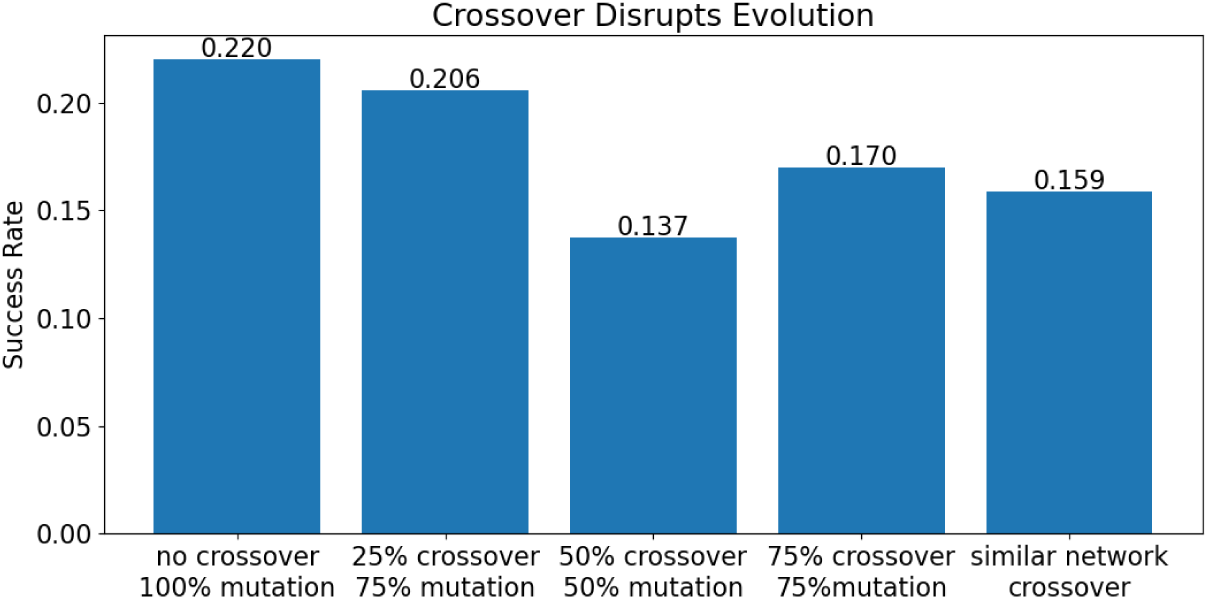
Evolution success rate of 700 trials with no crossover, 0.25 and 0.75 probability of crossover, 50-50 exclusive chance of crossover or mutation, and 0.75 probability of crossover with more lenient crossover method for networks with similar fitness.

With a target species of 10 (the default condition), evolution trials resulted in sustained oscillators at a rate of 0.22 (154/700). Without speciation, the success was significantly reduced (p *<*0.0001), with oscillators occurring at a rate of 0.12 (83/700). The batch with a target of 30 species also had a success rate of 0.22 (153/700).

Increasing the target number of species to 50 significantly (p=0.002) worsened results with a success rate of 0.16 (110/700). Similarly, decreasing the target number of species resulted in a success rate of 0.13 (89/700). This success is still a significant improvement over evolution without speciation (p *<*0.0001).

Speciation increases the success rate of evolution by sheltering innovations and maintaining population diversity. However, with too many species, there are not enough individuals in any given species to thoroughly explore the space and the success rate is reduced. Similarly, with too few species, there are not enough groups to more broadly explore the solution space and the success rate is reduced.

### 3.3 Crossover

To assess the effects of crossover on evolutionary success, batches of 700 evolution trials without crossover and with several type of crossover were compared. None of the crossover batches tested here had better evolutionary success than the batch without crossover.

First, the crossover technique described previously was compared to a batch with no crossover. Both batches had a target species of 10 and all other parameters in common. The batch with without crossover had 0% chance of crossover and 100% chance of mutation. 22% (154/700) of trials in this batch resulted in oscillators The batch with crossover used the crossover procedure described in the Methods section. The chance of crossover was 75% and the chance of mutation was 75%. All networks were either crossed over or mutated (*p_crossover_* + *p_mutation_* − *p_crossover_* ∩ *p_mutation_* = 1). This meant that networks had a 50% chance of being crossed over and then mutated (*p_crossover_* ∩ *p_mutation_* = *p_crossover_* + *p_mutation_* − 1). With these settings, evolution trials with crossover resulted in oscillators at a rate of 17% (119/700) of the time. This was significantly lower than the success rate without crossover (p = 0.001181).

It was possible that allowing some networks to undergo both crossover and mutation was too much change in a single step and was disrupting any progress towards a solution. In the next batch, networks were only allowed to undergo crossover or mutation (50% of each), but never both. In this case, 13.7% (96/700) trials resulted in oscillators, significantly less than both the batch without crossover and the batch with the original crossover method.

In a second attempt to address the problem of too much change, the chance of crossover was reduced from 75% to 25%. For this batch, 20.6% of trials resulted in oscillators. This is significantly better than the batch with a 75% chance of crossover (p=0.014), but not significantly different than the batch with no crossover.

With the original crossover method described in **??**, when two networks have the exact same fitness, one is randomly chosen as the more fit network. It is highly improbable that two networks would have exactly the same fitness, but this prompted the thought that there may be some benefit to more leniently combining two networks with similar fitness scores. For the next batch, networks with a fitness score within 5% of one another underwent a slightly different crossover method. Like the original crossover method, all reactions that were present in both parent networks were passed to the offspring, with the rate constant chosen randomly from one parent or the other. Reactions that were not shared between the two networks were passed down with a 0.5 probability. As with the original crossover method, reactions that were inactive had a 0.25 probability of reactivation. The success of this batch (16.1%, 111/700) was no different than the success of the original crossover batch and both were worse than the batch without crossover (p *<*0.0001).

### 3.4 Elitist Selection

Populating each generation begins with elitist selection. For each species, 10% of its allocated offspring are unmodified copies of the best networks from the previous generation. In cases where a species is only allocated a single offspring, a mutation is performed on the best network in the species. If the mutation improves the network, the network is passed on to the next generation with the mutation. If the mutation does not improve the network, the unmodified network is passed on to the next generation.

To explore the influence of elitist selection, batches of 700 trials of evolution were run with and without elitism. For batches with elitism, levels of 10, 25, and 50% were tested^1^. Eliminating elitism resulted in drastically reduced success rates, with only 1% (7/700) trials resulting in oscillators. However, in batches with elitism, the percentage of high fitness models copied to the next generation did not result in significantly different success rates. Batches with 10, 25, and 50% elitism had success rates of 0.220 (154/700), 0.224 (157/700), and 0.197 (138/700) respectively (figure 11). It is likely that evolution is driven primarily by the top few individuals. When elitism is removed, these top individuals are not preserved and evolution mostly fails. However keeping more individuals in addition to these top performers does not improve success rates.

**Fig 11.**
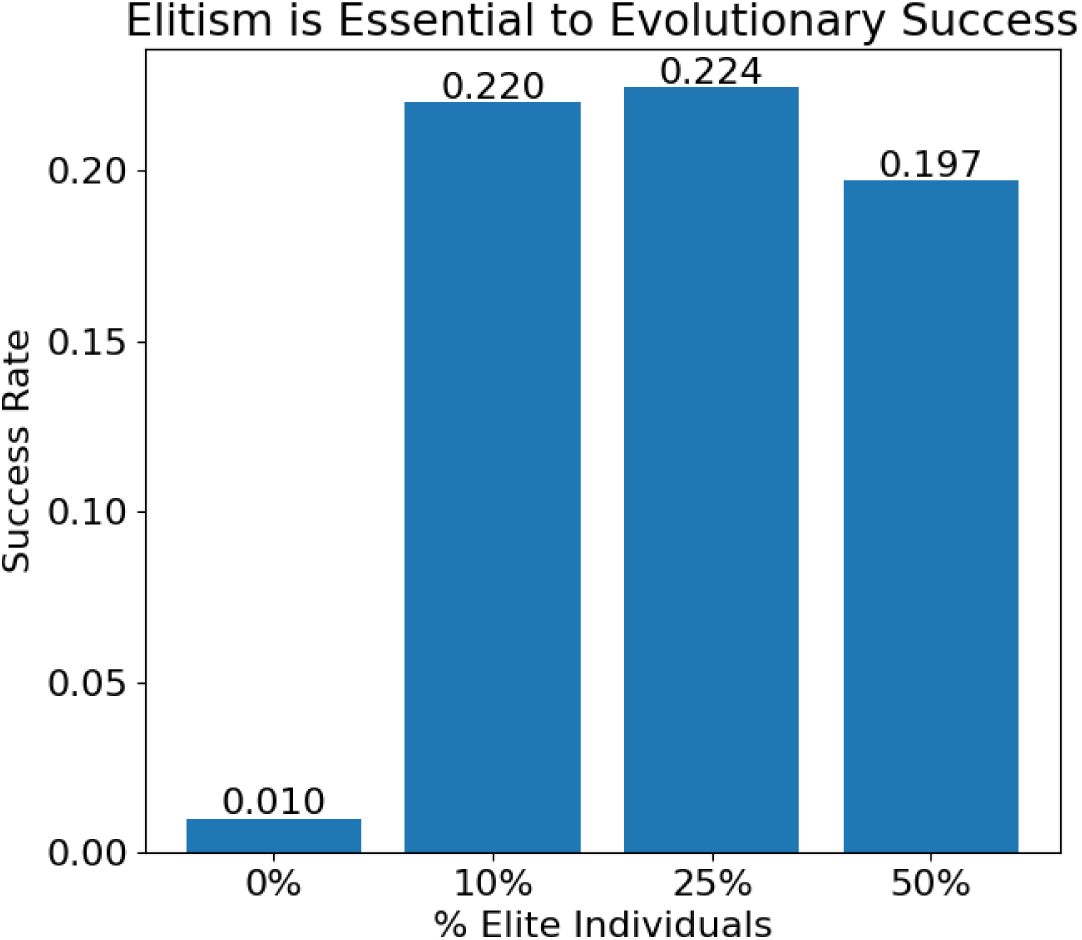
Evolution success rate of 700 trials with no speciation and with a target species of 10, 30, and 50.

**Fig 12.**
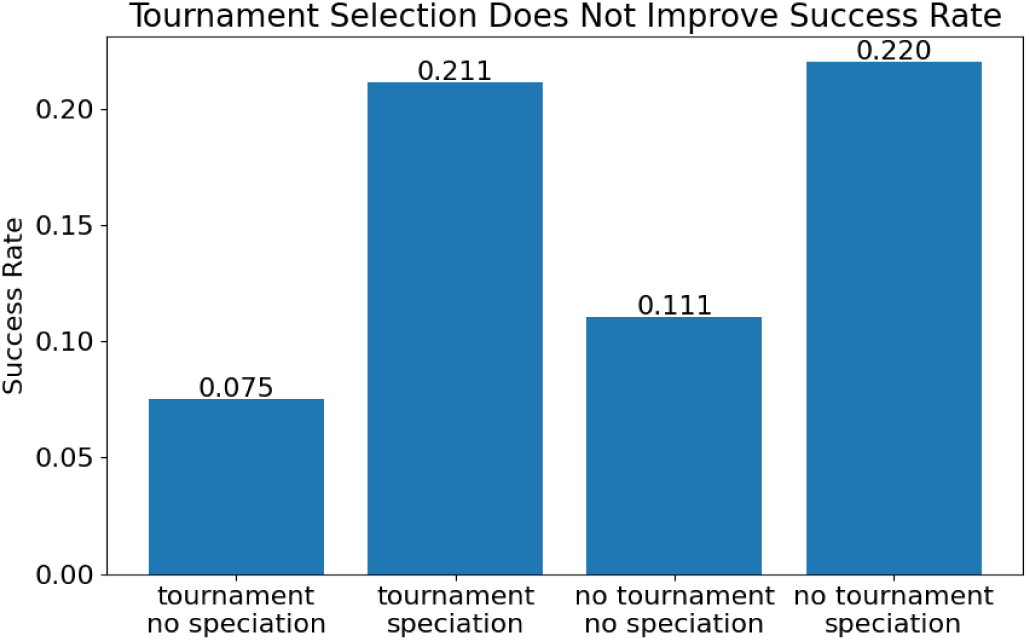
Evolution success rate with and without tournament selection and with and without speciation. Speciation influences success rate far more than tournament selection (n=1400 for each no speciation batch, n=700 for speciation batches).

### 3.5 Tournament Selection

After the elite networks were copied to the next generation, the remaining offspring were produced by randomly selecting networks (including the elites, excluding the bottom 10%) to mutate or crossover. As demonstrated in section 3.4, elitism is crucial to the algorithms success. But could a different selection strategy for the non-elite offspring improve success rate? To test this, tournament selection, with and without speciation, was used to select the non-elite offspring.

Tournament selection involves running several comparisons of a few individuals chosen at random from the population [25, 26]. The best of the subset is chosen for crossover or mutation. It is efficient to code and scales well population size and objective function. In this case, two individuals from the same species were randomly chosen (with replacement) and the individual with the better fitness score was allowed to mutate or crossover and continue to the next generation.

Tournament selection trials with a target species number of 10 and without speciation were compared to trials where non-elite networks were randomly selected. The trial with tournament selection and no speciation performed worst, with only 7.5% (105/1400) of trials resulting in an oscillator. Interestingly, random selection fared better, with 11.1% (155/1400) of trials resulting in oscillators (p *<*0.0001). This result was unexpected, as it would seem that biasing selection towards more fit networks with tournament selection should improve the results. However, it is possible that random selection helped maintain diversity by including less fit networks that ultimately developed into good solutions.

When considering tournament selection with speciation, results improved, with 21.1% (148/700) of trials resulting in an oscillator. However, this result in not significantly different than random selection with speciation, which resulted in oscillators in 22.0% of trials (154/700). This bolsters the conclusion that speciation drives evolutionary success more than any other factors explored here.

### 3.6 Increased Population size

Unlike other hyperparamaters discussed here, increasing the population size has a direct effect on computation time, as numerically solving the ordinary differential equations (ODE) is the rate limiting step of the algorithm. However, increasing the population size increases the number of individuals exploring the solution landscape and thus increases the chances of finding a good solution.

When the population size was doubled, from 100 individuals to 200 individuals, evolutionary success significantly improved from 22% to 26% (p = 0.012), with 182 of 700 trials producing an oscillator. However, this computation took roughly twice as long. The batch with 100 individuals resulted in 154 oscillators (out of 700 trials). Running this batch twice would result in 300 oscillators and take roughly the same amount of time as a 200 individual population that produces 180 oscillators.

Next, a batch with a population of 200 and a target of 20 species was run. For a population of 100, a target species of 10 was shown to result in the highest success rate, so this experiment sought to test if the optimal target number of species scaled with population size. This appeared to be the case as 38.5% (77/200) trials in this batch resulted in oscillators, a significant improvement over both 200 individuals with 10 species and 100 individuals with 10 species.

### 3.7 More Generations

Like population size, the number of generations directly effects computation time. However, increasing the number of generations allows more time for the algorithm to discover and optimize solutions and increase evolutionary success.

The maximum number of generations was doubled, from 800 to 1600, and evolution was run with default parameters: no crossover, 100 individuals, target species of 10. Trials in this batch resulted in oscillators 37% (74/200) of the time. The fitness of the top network in each generation was tracked and averaged across 200 trials (figure 13). After 1600 generations, the average top fitness continues to rise suggesting that the success rate may further increase with more generations. When the number of generations was doubled again, from 1600 to 3200, the success rate was 50% (100/200). At 3200 generations, the average fitness still did not plateau suggesting that a higher success rate may be possible at the expense of computing time.

**Fig 13.**
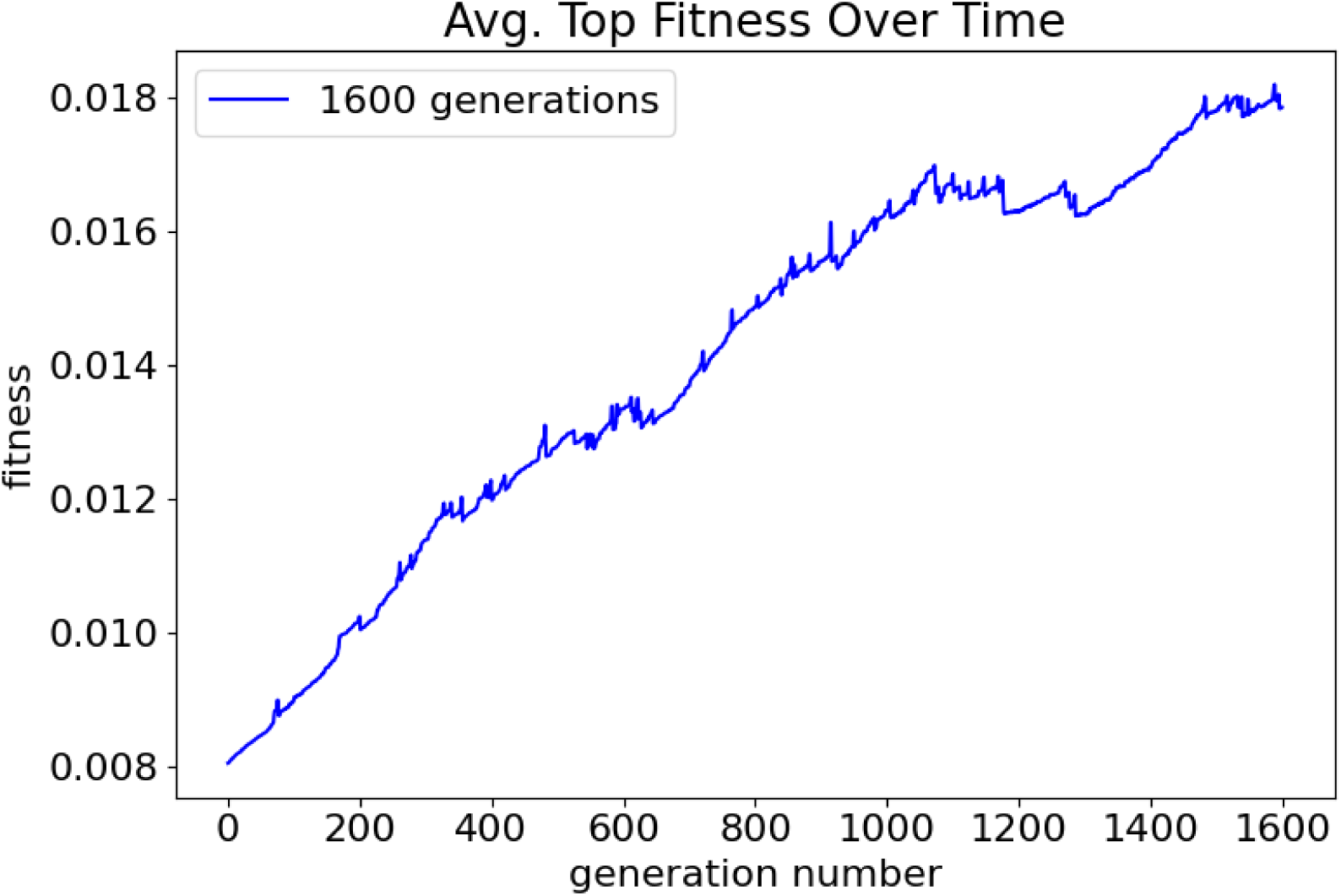
Average fitness of the best network of each generation over 200 trials. Small spikes can likely be attributed to writing out networks with high fitness and discontinuing evolution trials.

### 3.8 Evolving Oscillators with Optimal Settings

With the insights from these experiments, a batch of 720 trials was run with the best known hyperparameters. Two hundred individuals with a target of 20 species were evolved over 1600 generations without crossover and without tournament selection. Of the 720 trials, 374 resulted in oscillators for a success rate of 51.9%. It is likely that increasing the number of generations to 3200 or higher would further increase success rate, but at a cost of computing time.

Examples of time series data generated by randomly selected evolved oscillators are shown in figure 14. There are a variety of oscillation shapes, amplitudes, and frequencies demonstrating that there are many ways that the objective criteria can be met to produce an oscillator. This suggests that there are a variety of different oscillators in the evolved networks. The networks were not examined for duplicates.

**Fig 14.**
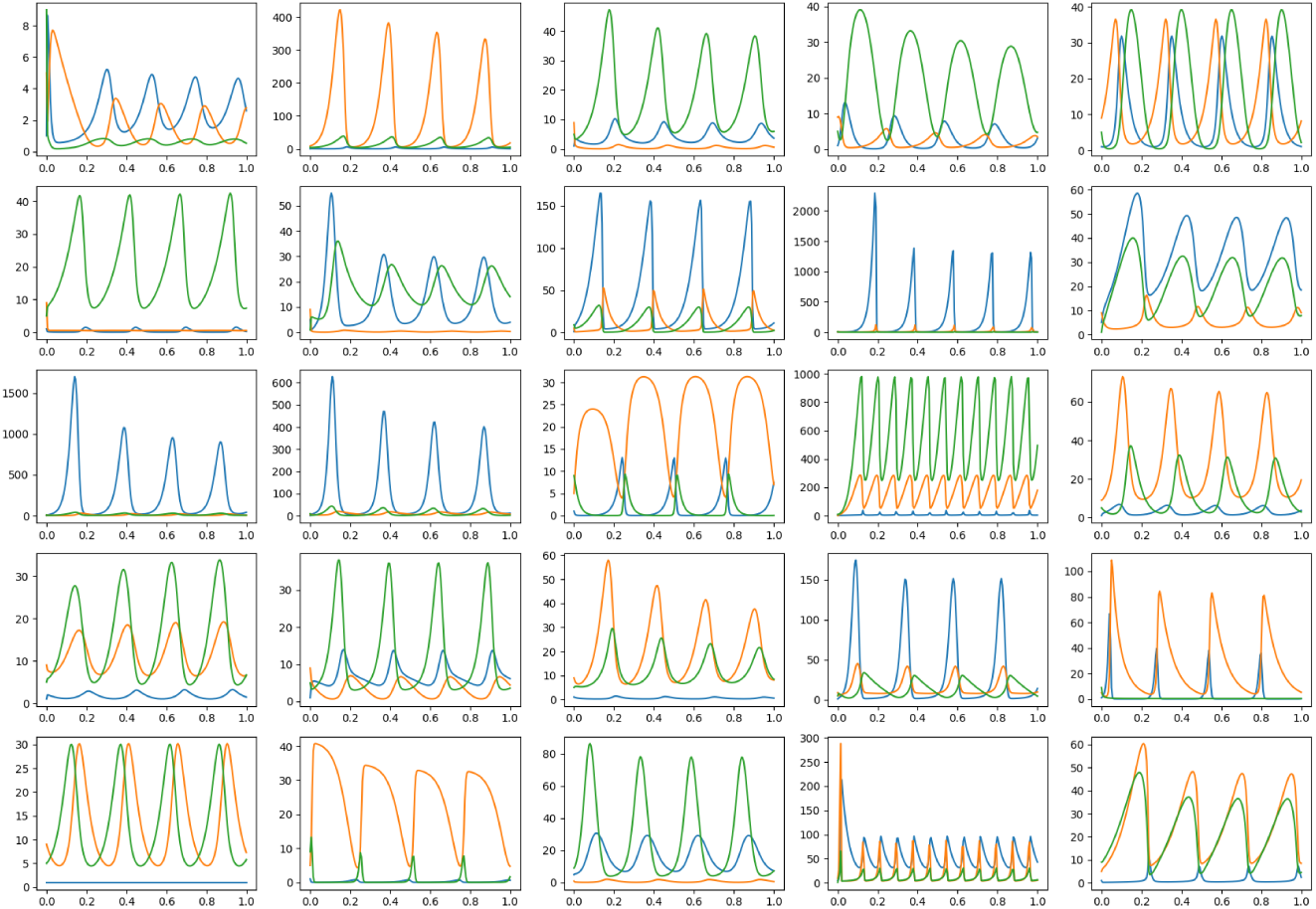
Time series data from an assortment of evolved oscillators.

Examples of three evolved networks and their time series data are shown in figure 15. Each network has been processed to remove all reactions that are not necessary for oscillation. The three networks vary in complexity and produce different oscillations in terms of frequency, amplitude, and shape. Despite the difference in complexity, the second and third networks generate a similarly shaped time series. Interestingly, the third network has completely ignored the third chemical species and recreated the Lotka-Volterra predator-prey model.

**Fig 15.**
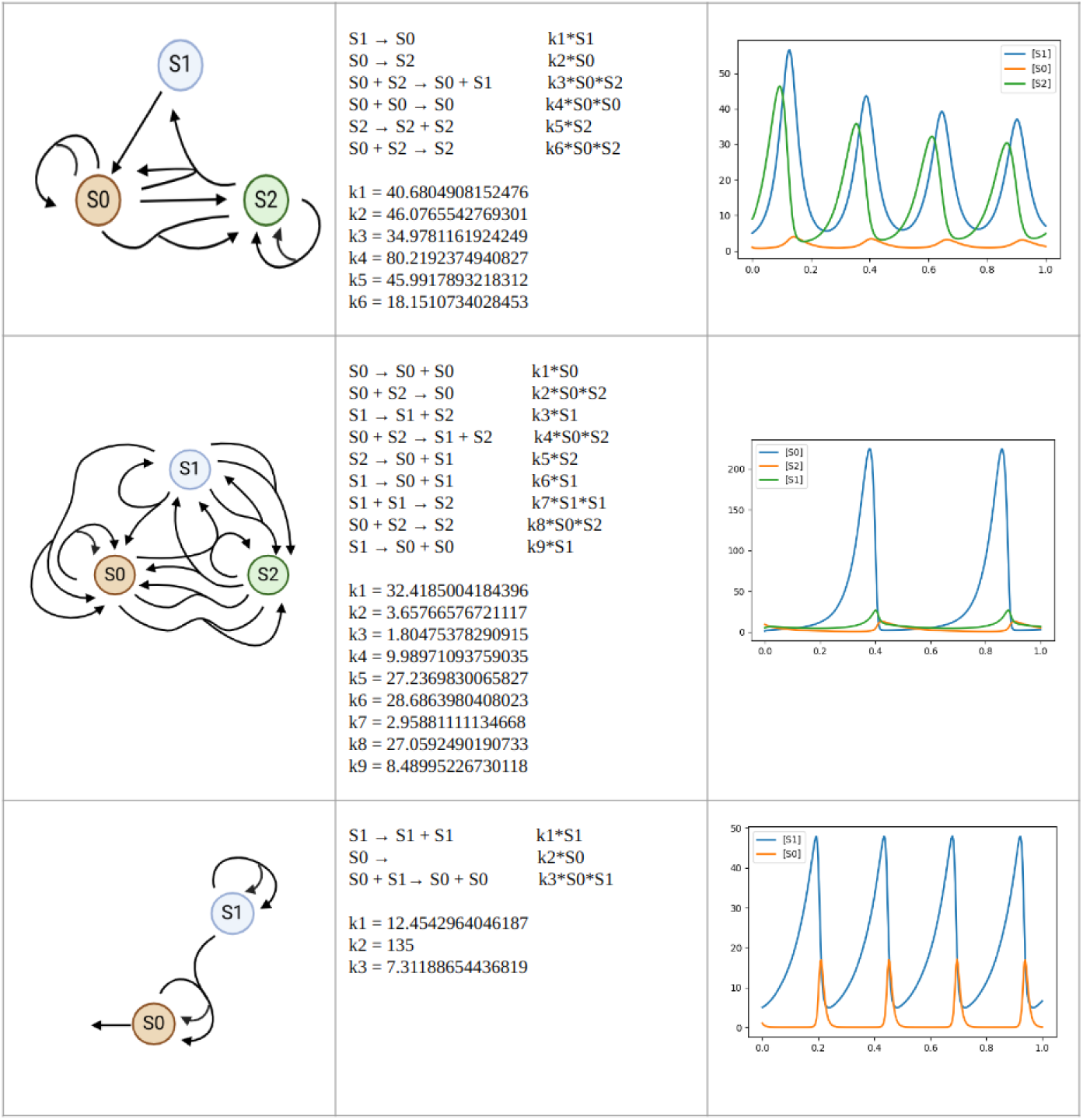
Examples of oscillatory networks and their time series. The bottom network is the Lotka-Volterra predator-prey network.

The top fitness values at each generation for five successful evolution trials are shown in Figure 16. Some networks, such as those shown in blue and purple, make a series of mutations that rapidly improve the fitness. Others develop more gradually. Trials with a fitness value of 0.05 or higher were stopped and saved, but 0.05 is a very conservative cutoff. It is likely that a cutoff value of 0.01 would capture most if not all oscillators.

**Fig 16.**
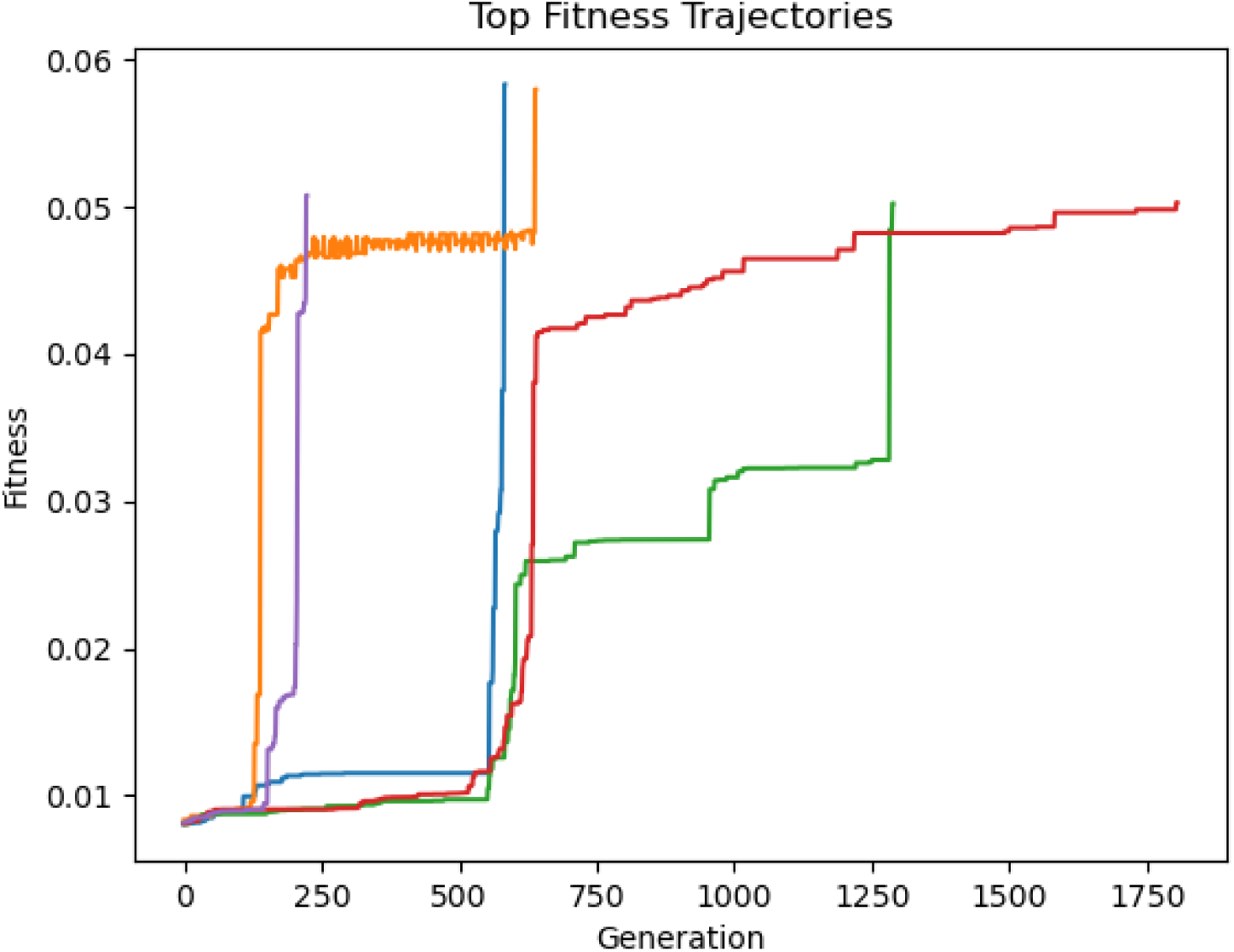
Examples of fitness trajectories of successful evolution trials.

### 3.9 Fitting Time Series Data

Beyond generating oscillators, the ReactionNetworkEvolution.jl software is intended to implement an evolutionary algorithm for more broad systems biology problems. One application could be evolving reaction networks that match time series data. To test if ReactionNetworkEvolution.jl could generate networks to match time series data, a simple, 3 chemical, 5 reaction synthetic network was constructed and its time series data collected (Figure 17). Then the ReactionNetworkEvolution.jl software was given the time series data as an objective.

**Fig 17.**
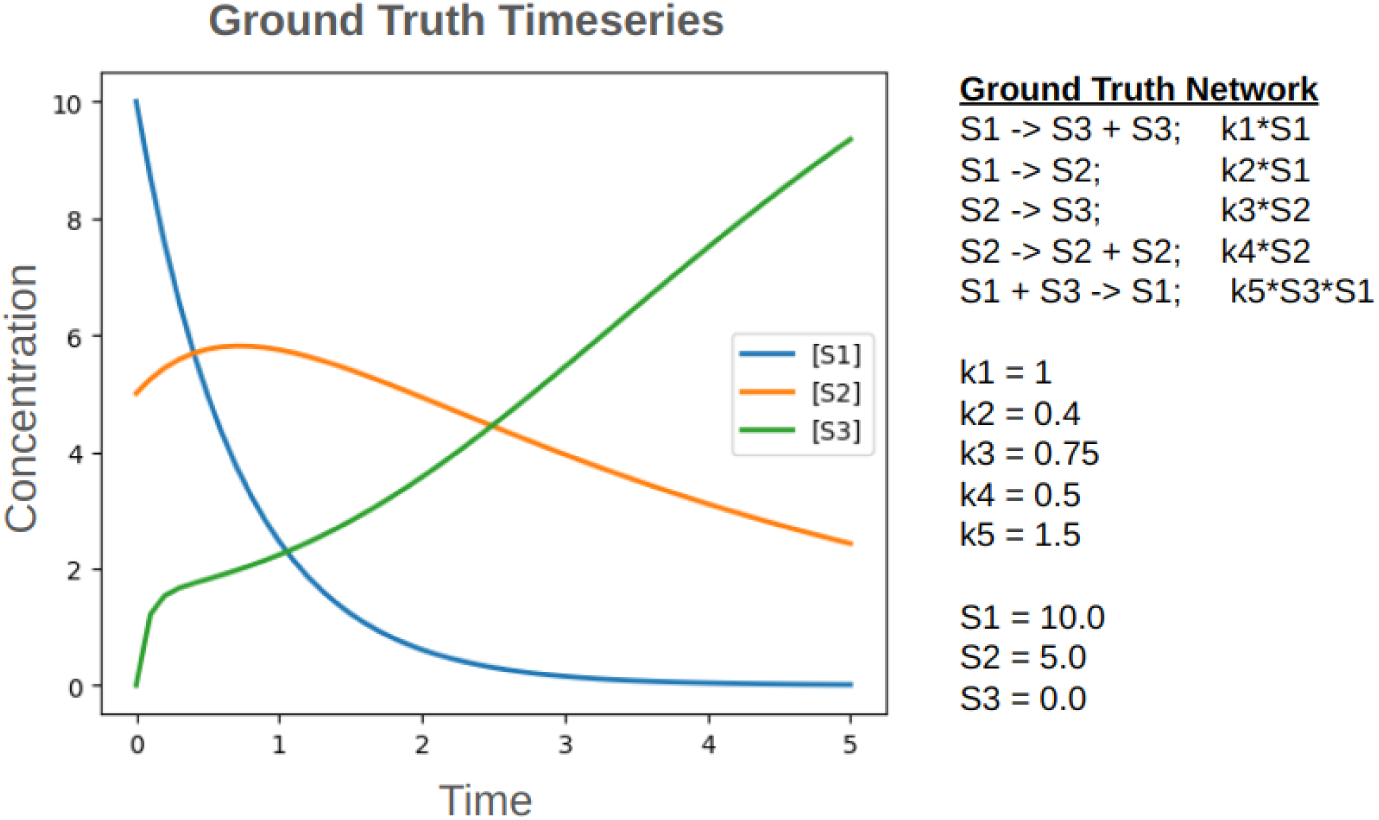
A synthetic network and its time series data.

Four hundred trials of evolution were run, each with 1600 generations in an effort to match the ground truth time series data. The time series data of the 12 best evolved networks all closely match the dynamics in the ground truth network (Figure 18). The best matching time series (top row left of Figure 18) is shown overlaid with the ground truth time series in Figure 19. The solid lines are data from the ground truth network and the dashed lines are from the evolved network.

**Fig 18.**
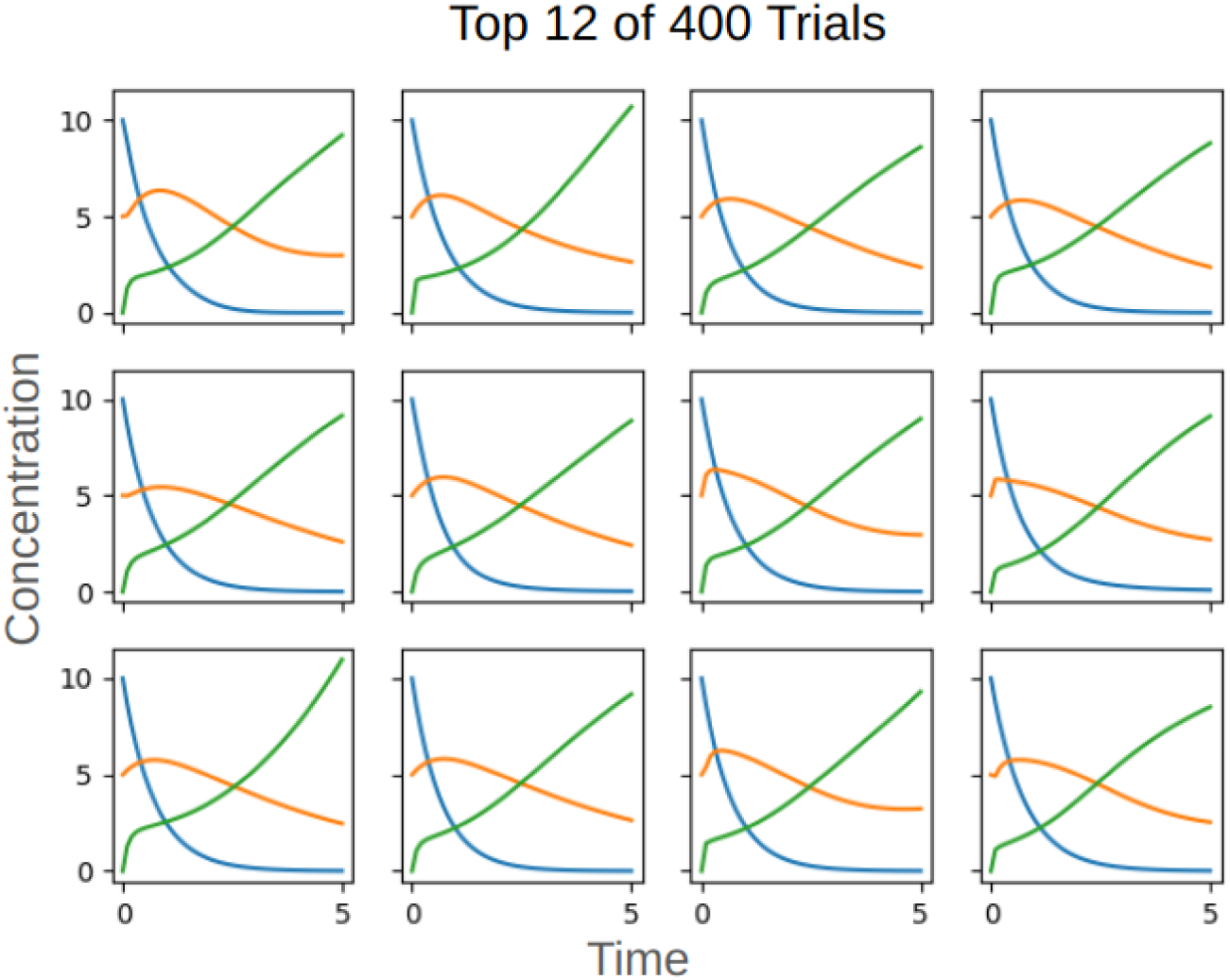
The 12 trials that most closely matched the time series data.

**Fig 19.**
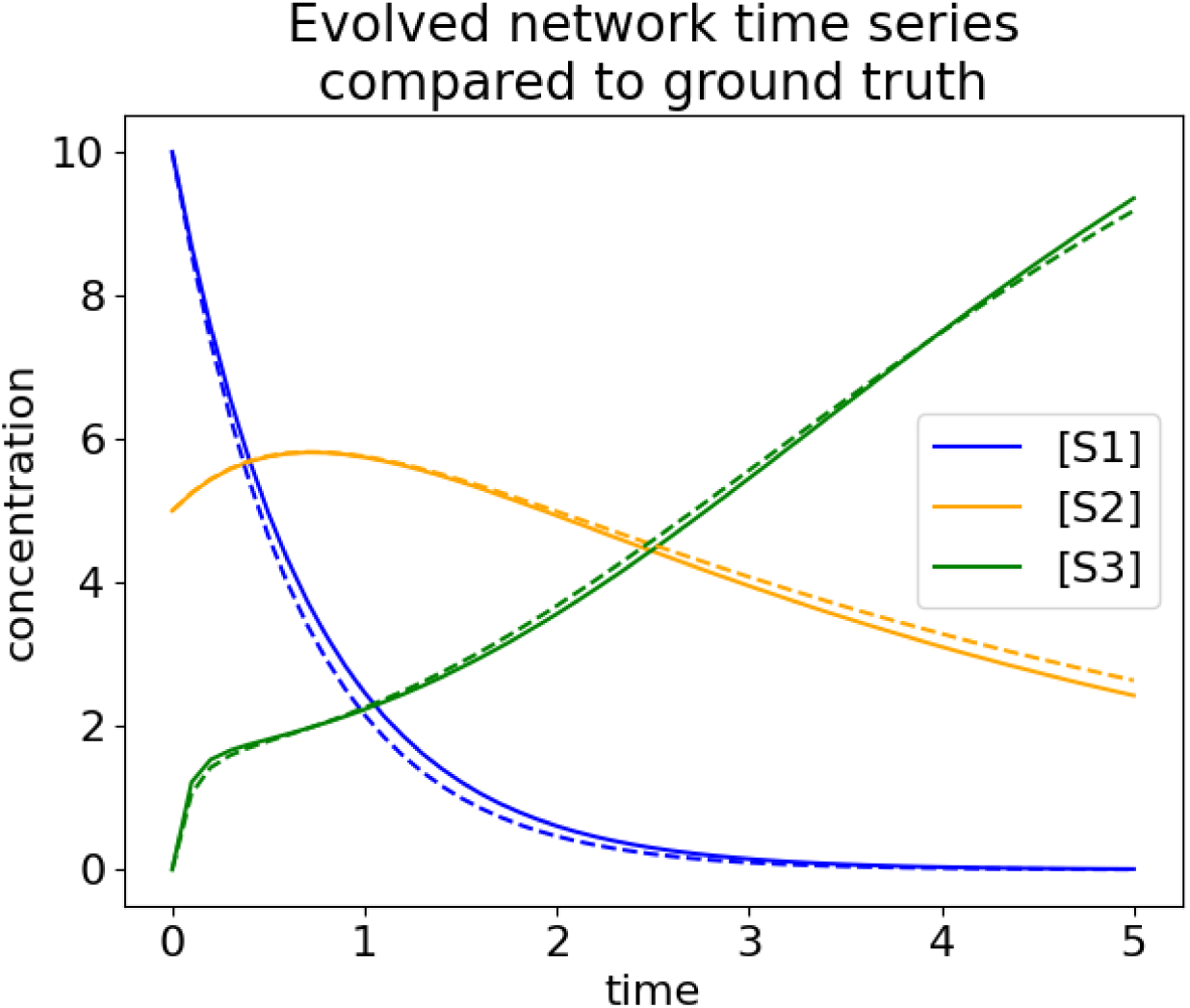
Time series of the ground truth network (solid lines) compared to the best evolved network (dashed lines).

Interestingly, the evolved network that most closely matched the ground truth time series data only had one reaction in common with the original network (Figure 20). The last reaction, S1 + S3 → S1, occurred in 10 of the 12 best networks. The next most common was the reaction S2 → S3, which occurred in 7 of the 12 best networks. All networks that had the reaction S2 → S3 also had the reaction S1 + S3 → S1.

**Fig 20.**
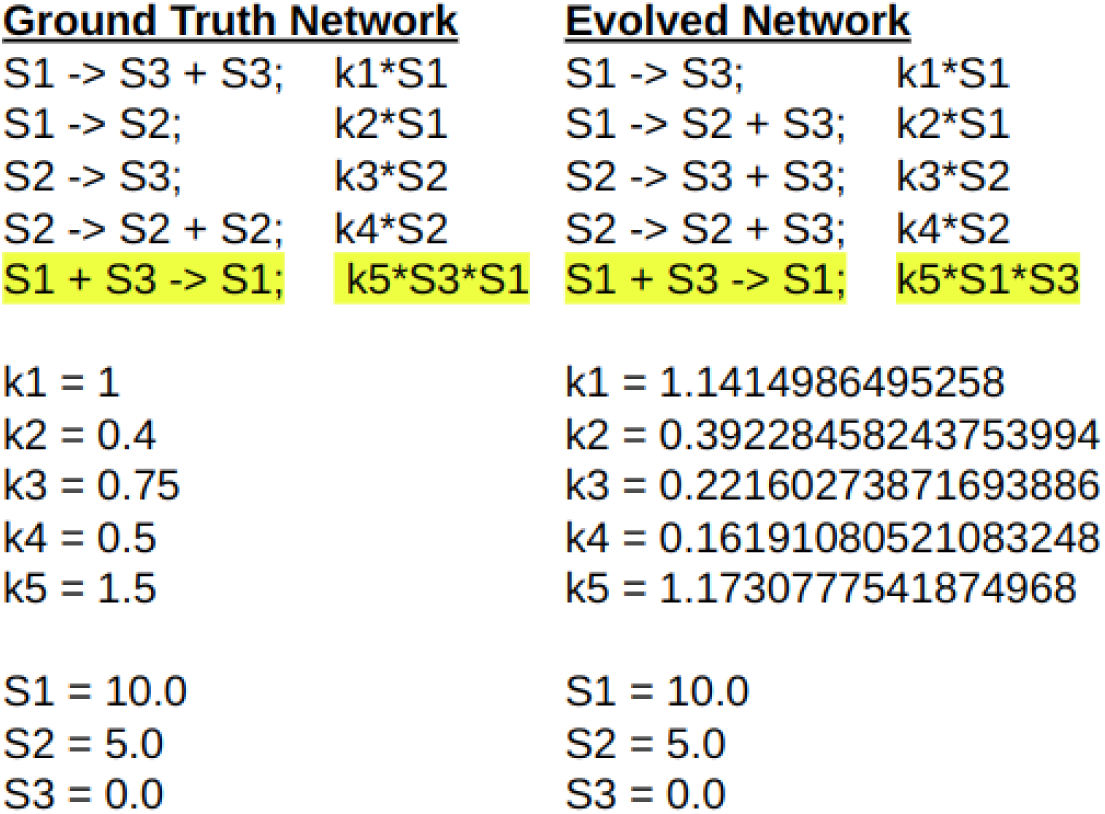
The ground truth network compared to the best evolved network

## 4 Discussion

Speciation was shown to significantly benefit evolutionary success by protecting innovations and maintaining evolutionary diversity throughout the process. Fine tuning is necessary to achieve the maximum benefit of speciation. The target number of species must be chosen with consideration of the population size. Additionally, *δ* and *δ* step sizes may require adjustment for different population sizes, target number of species, and different objectives.

Elitism is also essential to evolution as it preserves the best individuals of each generation. It pairs well with speciation as it allows the best individuals not just from the whole population but from the each species to be passed on to the next generation. Interestingly, the evolutionary success did not seem to vary with different levels of elitism suggesting that evolutionary progress is driven primarily by the top individuals of each species and that including additional elite networks does not improve success. This seems to be confirmed by the observation that alternative selection strategies for non-elite individuals did not affect evolutionary performance.

Although crossover has shown success in other topological evolution problems [16, 17], this was not the case with mass-action networks. Both artificial neural networks (ANNs) and genetic regulatory networks (GRNs) (domains where crossover has been shown effective) have more edges that can more easily be separated from their weights. In both cases, edges simply connect two nodes together. For ANNs, all edges activate the output node and have a weight which increases or decreases the signal strength. GRNs are similar but edges can also be inhibitory. The weights of the edges are not influence by the nodes the edge is connected to. On the other hand, in this work, reactions are analogous to edges and can connect two to four nodes (chemical species). The rate constants are analogous to weights. But unlike weights, the extent to which one node activates another (or the speed at which a reactant becomes a product) is also influenced by the reactant’s concentration. That is to say, in the case of mass-action chemical reaction networks, it is far more difficult to isolate subsets of a network and maintain their function. When crossover occurs, it tends to disrupt the finely tuned developing solutions by changing several rate constants simultaneously. This could explain why crossover reduces success in this application despite its utility in other domains.

Although this algorithm is faster than previous iterations, it still computationally expensive. A single trial of 800 generations and 100 individuals take approximately 90 seconds on average. Given the success rate using these settings, this amounts to approximately 3 minutes per oscillator per CPU. The vast majority of this time is consumed by numerically solving the ODEs. The software uses a custom function to compute the rate of change of each concentration, but the solver is from julia’s DifferentialEquations and Sundials libraries, both highly optimized for performance. Additionally, ReactionNetworkEvolution.jl uses strict typing, pre-allocation wherever possible, and hash maps to further improve performance. Although the current implementation of ReactionNetworkEvolution.jl does not make use of parallelization within a single trial, in practice multiple trials can be dispatch to different CPUs using a bash script or job array. Future efforts to improve the success rate are likely to be more fruitful than efforts to increase computational performance.

Using an adaptive speciation threshold, *δ*, did not directly influence evolutionary success, but it maintained the target number of species which did influence success. Without an adaptive *δ*, the user would have to manually tune *δ* through trial and error to achieve the target number of species. Given the utility of this adaptive feature, it may be useful to implement a similar strategy for other evolutionary hyperparameters. For example, the probability of changing the topology (adding or deleting reactions) versus changing rate constants remains the same throughout the evolutionary process. Early on, it may be useful to change topology more frequently in an effort to find a decent solution. However, later in the process, when a promising solution has been found, it is more useful to modify rate constants in an effort to optimize the solution. Changing topology may be wasted effort and may divert resources that may be better spent optimizing an existing solution. In future iterations of ReactionNetworkEvolution.jl, it may be useful to change the balance of topological versus rate constant mutation over time or as fitness reaches a pre-specified threshold.

Similarly, as evolution progresses and homes in a promising solution, smaller rate constant changes may be necessary to explore a small area in the solution space in more detail. The amount by which rate constants are changed could be adaptively reduced over time, either based on fitness or generation number.

Another means of more thoroughly exploring a promising solution space as evolution progresses could be altering the metric used to determine the “distance” between two networks, and thus their species. In the current iteration of ReactionNetworkEvolution.jl, the distance between two networks is entirely determined by the amount of reactions that the two networks do not share with no consideration to the rate constants for shared reactions. Networks with the same topology are considered identical even if rate constants for their different reactions vary by orders of magnitude. Including the differences in rate constants when considering distance could improve evolution by encouraging more thorough explorations of viable solution spaces towards the end of the process. As networks become more topologically similar as evolution progresses, speciation would be driven by differences in rate constants instead of topology. This would direct more resources towards optimizing a decent solution as opposed to continuing to explore the broader solution space.

One flaw of the existing iteration of ReactionNetworkEvolution.jl is the tendency to add extraneous reactions. It appears that more often than not, deleting a reaction reduces fitness whereas adding a reaction may slightly improve fitness. Although there was equal probability of adding and deleting reactions, networks at the end of evolution tended to be much larger (approximately 15 reactions) than at the start (5) reactions. In some cases, this may not be of concern. For example, if the objective is simply to create a population of oscillators, it may not matter that some of them are overly complicated. However, if attempting to gather simple networks to fit time series data, the tendency to over complicate networks could be detrimental. Overfitting by over complicating models (for example, by including too many predictors) is a well-known problem in machine learning. Often, the solution is to penalize machine learning models that use too many features. A similar technique could be implemented here, penalizing candidate networks for each additional reaction. Penalization was briefly explored, but was found to reduce the success rate. Further fine tuning of a fitness function that addresses network complexity may increase the success rate and result in less complex models.

Isomorphism was not considered during of after the evolution process. Networks were only considered similar or identical if both their reactions and the names of the chemical species involved in the reactions matched. However, given the arbitrary naming of the chemical species, it is possible to have two identical networks but with shuffled chemical species names. During the evolution process, this might result in two or more species groups effectively populated with the same network. Similarly, after multiple trials of evolution, there may be a number of oscillators that are isomorphic.

The issue of isomorphism could be partially addressed in future versions of ReactionNetworkEvolution.jl. During evolution, the names of chemical species could be normalized during fitness evaluation such that the chemical species that most closely matched the input time series was named “*S*_0_,” followed by “*S*_1_,” and so on. One possible problem with this proposal is that different reaction rates may cause different chemical species to best match the input time series data despite two networks being topologically identical.

Identifying isomorphic duplicates after multiple evolution trials could be addressed with multiple pairwise comparisons. For each pair of networks with 3 chemical species, six comparisons between the two networks would be sufficient to identify if two networks are the same despite having shuffled chemical species names. For a population of several oscillators, this would amount to 6 *^n^* comparisons, where n is the number of oscillating networks to be evaluated. A population of 40 oscillators would require 4,680 comparisons to identify all duplicate networks.

Additionally, the algorithm has demonstrated its ability to produce oscillators, but its ability to match other network behaviors is untested. It successfully generated a network to match simple time series data, but did not manage to recover the network that produced the data. This is expected given the limited amount of information about the network provided to the software. More testing and algorithm development could improve ReactionNetworkEvolution.jl’s performance in tasks beyond generating oscillators.

## 5 Conclusion

The NEAT algorithm for evolving artificial neural networks was successfully adapted to mass-action chemical reaction networks. The experimental results showed that speciation and elite selection drive evolutionary success. Unlike other domains, crossover negatively impacts evolution of oscillatory mass-action networks.

The adapted evolutionary algorithm is packaged in a julia module, ReactionNetworkEvolution.jl, for easy use and customization. ReactionNetworkEvolution.jl boasts at least a four-fold improvement in success rates over previous iterations and has considerably better computational performance. Using the default settings, oscillators are generated every three minutes on average using a single CPU.

Future work will explore the evolution of more complicated network behaviors. Other directions include improving the algorithm’s ability to match time series data when given more information about the network or to suggest what additional information is necessary to narrow down the set of networks that can recapitulate the input time series data.

## Acknowledgments

This work was partly supported by NIH Biomedical Imaging and Bioengineering award P41 EB023830. The content expressed here is solely the responsibility of the authors and does not necessarily represent the official views of the National Institutes of Health, or the University of Washington. LT thanks Joe Hellerstein for useful discussions.

## Footnotes

1 This is the percentage of the *next* generation that is composed of unmodified, higher fitness models. For example, if a species has 20 members but is only allowed 10 offspring with an elitism rate of 10%, then the single best network of the 20 will be passed down to the next generation unmodified.

